# Fitting mathematical models of biochemical pathways to steady state perturbation response data without simulating perturbation experiments

**DOI:** 10.1101/183194

**Authors:** Tapesh Santra

## Abstract

A common experimental approach for studying signal transduction networks (STNs) is to measure the steady state concentrations of their components following perturbations to individual components. Such data is frequently used to reconstruct topological models of STNs, but, are rarely used for calibrating kinetic models of these networks. This is because, existing calibration algorithms operate by assigning different sets of values to the parameters of the kinetic models, and for each set of values simulating all perturbations performed in the biochemical experiments. This process is highly computation intensive and may be infeasible when molecular level information of the perturbation experiments is unavailable. Here, I propose an algorithm which can calibrate ordinary differential equation (ODE) based kinetic models of STNs using steady-state perturbation responses (SSPRs) without simulating perturbation experiments. The proposed algorithm uses modular response analysis (MRA) to calculate the scaled Jacobian matrix of the ODE model of an STN using SSPR data. The model parameters are then calibrated to fit the scaled Jacobian matrix calculated in the above step. This procedure does not require simulating the perturbation experiments. Therefore, it is significantly less computation intensive than existing algorithms and can be implemented without molecular level knowledge of the mechanism of perturbations. It is also parallelizable, i.e. can explore multiple sets of parameter values simultaneously, and therefore is scalable. The capabilities and shortcomings of the proposed algorithm are demonstrated using both simulated and real perturbation responses of Mitogen Activated Protein Kinase (MAPK) STN.

**Availability:** All source codes and data needed to replicate the results in this manuscript are available from https://github.com/SBIUCD/MRA_SMC_ABC1

## Introduction

Steady state perturbation responses (SSPRs) of STNs are quantified by (a) perturbing the individual components of the STN using chemical inhibitors, siRNAs, viral vectors or plasmids; (b) following each perturbation letting all components of the STN to relax into a steady state; and (c) subsequently measuring the phosphorylation levels of each component (*1-5*). The effects of each perturbation spread through the network of biochemical interactions and influence the activities of different components to various extents. Such data contains information about the causal relationships between different components of the STN, and is typically used to reconstruct its topology (*1-8*). The topological models shed some light on patterns of interactions between biochemical molecules, but lack sufficient mechanistic details for predicting how the STN will respond to external stimuli and/or interventions. More detailed mathematical models of STNs can serve this purpose (*9*). Such models can be developed using the reconstructed topological models as guides for determining which interactions do and do not take place in the STN (*1, 10*). However,these detailed models are parameter rich and calibrating these parameters is challenging. Existing calibration algorithms iteratively explore different sets of parameter values (*1, 11-15*). In each iteration, a random or semi-random set of values are assigned to the model parameters, the model is then used to simulate the same perturbations as was performed in biochemical experiments, the simulated SSPRs are then compared with the experimental data. The sets of parameters which provide an acceptable level of similarities between simulated and experimental SSPRs are then used as calibrated parameter values (*1, 11-15*).

However, simulating several model perturbations in each iteration is computation intensive.Additionally, such simulation requires detailed knowledge of the way the perturbations were performed and their effects on the STNs. This information is often not readily available. Even when such knowledge is available it is not straightforward to simulate SSPRs. For instance, biochemical STNs are often perturbed using inhibitors. Even if the type and concentration of the inhibitor is known, simulating perturbations mediated by the inhibitor requires one to incorporate additional equations describing the interactions between the inhibitor and its target into the mathematical model. This increases the number of unknown parameters that needs to be calibrated and thereby reduces model identifiability. Here, I propose a calibration algorithm which avoids simulating perturbation experiments altogether when fitting models to SSPR data. The algorithm calculates a scaled Jacobian matrix of the mathematical model of the STN from SSPR data using MRA. It then calibrates the model parameters using an adaptive weight approximate Bayesian computation scheme (*16*) so that the scaled Jacobian of the calibrated model fits that calculated from the SSPR data. In the following sections we describe the details of this algorithm and demonstrate its applicability using simulated and real SSPRs of the MAPK STN.

## Method

### Bridging Jacobian matrix of ODE model with SSPR data using MRA

Let us assume that an STN contains *N* nodes which regulate each other’s concentrations. A mathematical model (*M***_*x*_**) that formulates how the interactions between the different nodes influence their concentrations consists of a set of ordinary differential equations (ODE) of the form 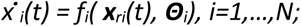 where 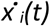 represent the rate at which the concentration (*x* _*i*_*(t)*) of the *i*^*th*^ node changes with time (*t*), *f*_*i*_ is a continuous function, ***x***_*ri*_*(t)* are the concentrations of the regulators of node *i* including itself, ***Θ***_*i*_ are the parameters of the function *f*_*i*_. The values of the parameters (***Θ****=*{ ***Θ***_*i*_ *,i=1,…,N*}) are unknown and needs to be estimated from experimentally observed data. The experimental data is generated by perturbing each node of the network, one at a time. Following a perturbation (*p*_*i*_) to each node (*i*), the STN is allowed to relax into a steady-state and the changes in the concentrations of all nodes (**x***=*{*x*_*i*_ *, i=1,…,N*}) in response to each perturbation (*p*_*i*_) are measured. Our objective is to use this data to fit the parameters (***Θ****=*{***Θ***_*i*_ *,i=1,…,N*}) of the mathematical model (*M***_*x*_**) of the STN without simulating the perturbation experiments during the fitting process. To do so, we exploit a relationship between the Jacobian matrix (***J****(t)*) of the ODE model and the experimentally observed SSPRs of the STN.

Note that at steady state 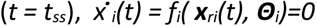 and therefore *df*_*i*_( ***x***_*ri*_*(t),* ***Θ***_*i*_*)=0.* Using chain rule of derivative

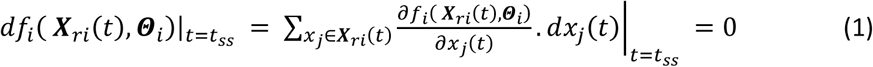

If we assume a hypothetical scenario where all but the i^th^ and j^jh^ nodes of the STN are kept fixed then the above equation reduces to:

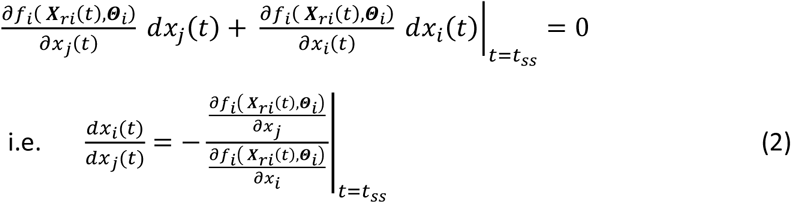

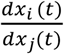quantifies the change in the concentration (*x*_i_) of node *i,* due to an infinitesimally small perturbation to node *j,* when the concentrations of all other nodes are kept fixed. In other words,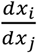 represents the influence of an infinitesimally small perturbation to node *j* on the concentration of node *i* when all interactions but the regulation of node *i* by *j* are disconnected. The numerator 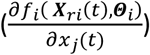and denominator 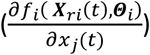 of the right hand side of Eq. 2 are in fact (i,i)^th^and (i,j)^th^ elements (***J_ii_, J****_ij_*) of the STN’s Jacobian Matrix (***J****(t)*) which is defined as **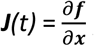** where ***f****=*{ *f (* ***x***_*ri*_*(t),* ***Θ***_*i*_*),i=1…N* } and ***x*** *=*{*x*_*j*_ *(t),j=1…N* }. Scaling both sides of Eq. 3 by the ratio (*x*_*j*_ *(t*_*ss*_*)/x*_*i*_*(t*_*ss*_)) of the steady state concentrations of nodes *j & i* gives us

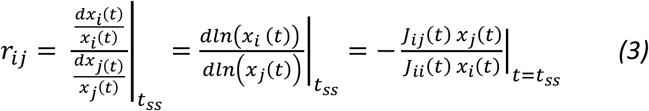

*r*_*ij*_ is known as the local response coefficient (LRC) of the regulation of node *i* by *j* and represents the change in the logarithmic steady state concentration of node *i* due to a small perturbation to node *j,* when all other nodes are disconnected. It was shown by Kholodenko et. al. (*17*) that the local response matrix (***r****=* {*r*_*ij*_*, i,j=1,…,N*} for notational convenience, can be calculated from the experimentally observed SSPRs by solving the following linear equations

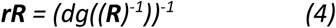

]Where ***R*** = {*R*_*ij*_ *, i,j=1,…,N* } is the global response matrix whose elements *R*_*ij*_ *= (Δln(x*_*i*_*)/ Δln(x*_*j*_)) are known as global response coefficients which represent the ‘global change’ (i.e. when the perturbation is allowed to propagate through the network) in the logarithmic concentration of node *i* due to an infinitesimally small perturbation to node *j*. Experimental perturbations are never infinitesimally small, therefore *R*_*ij*_ is calculated in an approximate sense from experimental SSPR data using the following formula (*17*):

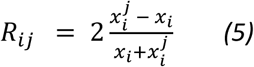

Here *x*_*i*_ and 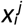 is the experimentally measured steady state concentrations of node *i* prior to and following a perturbation to node respectively.

Eqs. 3 allows us to calculate the local response matrix (**r**) of and STN using its ODE models without explicitly simulating the perturbation experiments. Therefore, estimating the model parameters (***Θ****=*{ ***Θ***_*i*_ *,i=1,…,N*}) boils down to the following two steps:

- **Step1:** Calculate the local response matrix ***r*** from experimentally observed SSPRs using Eqs. 5 and 4
- **Step2:** Simulate the local response matrix (***r***_*s*_) using Eqs. 3 for different values of model parameters and chose the sets of values which provide close match between ***r*** and ***r***_*s*_.

Searching for parameter values that minimizes the difference between ***r*** and ***r***_s_ out of infinitesimally many possibilities is a challenging task. There are several algorithms which provides efficient ways for exploring the parameter space (possible values of parameters). The algorithms relying on Approximate Bayesian Computation using Sequential Monte Carlo (ABC-SMC) (*15, 16*) are particularly attractive choice since they stem from the traditional Bayesian algorithms which allow one to incorporate prior knowledge of parameter values into the inference system, while making it possible to perform the search across multiple computers in parallel. Here we resort to a improved version of the original ABC-SMC (*15*) known as Adaptive Weight ABC-SMC (*16*). The algorithm is briefly described below including how we parallelize it.

### Exploring the parameter space using Adaptive Weight ABC-SMC

ABC is inspired by Bayesian Statistics and relies on Bayes principles which provides a framework for updating prior knowledge about an unknown variable or quantity using observed data. The prior and updated knowledge are represented in the form of probability distributions, known as prior and posterior distributions respectively, which formulates our initial guesses and updated estimates about the potential values of the unknown variables. Adaptive Weight ABC-SMC algorithm starts by assigning prior distributions *(P(****Θ***)) to the model parameters (***Θ***), and initializing a monotonically decreasing set of error thresholds (*ε*_*t*_*, t=1,…,T, ε*_*1*_*> ε*_*2*_*>…> ε*_*T*_) which will be used to refine the posterior distribution (***P***(***Θ****|****D*_*obs*_***), (****D*_*obs*_** is the observed data, which, in our case, is the local response matrix, i.e. **D**_obs_ = ***r***) of the model parameters (***Θ***) in a stepwise manner as described below (for details see (*15, 16*)).

**Step 1:** In the first step (*t=1*), a number of potential values (*Θ***^*1k*^**, *k=1,….)* of the model parameters (***Θ***) are sampled from their prior distributions (*P(****Θ***)). For each set of values (*Θ***^*1k*^**) a local response matrix (***r***_*s*_^*1k*^) is simulated using the ODE model (*M*_*x*_), and the error between the simulated (***r***_*s*_^*1k*^) and the SSPR derived (***r***) local response matrices are calculated using a distance measure (*d(****r***,***r***_*s*_^*1k*^)). If the error (*d(****r***,***r***_*s*_^*1k*^)) is less than the error threshold *ε*_*1*_, i.e. *d(****r***,***r***_*s*_^*1k*^*)< ε*_*1*_, then corresponding parameter values (*Θ***^*1k*^**) are kept for next iteration, otherwise discarded. This process is repeated until a desired (*N*_*ABC*_) number of parameters are kept (***Θ*^*1*^*=*** {*Θ***^*1n*^,** *n=1,..,N*_*ABC*_ }). Each of the selected values (*Θ***^*1n*^**) is assigned a weight (*ω*^*1n*^ *=1/N*_*abc*_).

**Step 2:** In the next step (*t=2*), one (*Θ***^*1n*^**) of the parameter values (***Θ*^*1*^**) which were not discarded in the previous step is selected with probability *p*^*1n*^ *ω*^*1n*^ *K*_*c*_(***r***_*s1n*_,***r***) where *K*_*c*_(***r***_*s1n*_,***r***) measures the closeness between ***r***_*s1n*_ and ***r***. A new parameter value (*Θ***^*2k*^**) is then proposed by sampling a proposal distributions (*P(Θ*^*2*^*| Θ*^*1n*^ )) which is conditioned on the selected value (*Θ***^*1n*^**). A local response matrix (***r***_*s*_^*2k*^) is then simulated using this parameter value (*Θ***^*2k*^**) and the error (*d(****r***,***r***_*s*_^*2k*^)) between the simulated (***r***_*s*_^*2k*^) and the SSPR derived (***r***) local response matrices is calculated. If the error (*d(****r***,***r***_*s*_^*2k*^)) is less than the error threshold *ε*_*2*_, i.e. *d(****r***,***r***_*s*_^*2k*^*)< ε*_*2*_, then newly sampled value (*Θ***^*2k*^**) is kept, otherwise discarded. This process is repeated until a desired (*N*_*ABC*_) number of parameters are kept (***Θ*^*2*^*=*** {*Θ***^*2k*^,** *k=1,..,N*_*ABC*_ }). The weights of the selected values are updated as follows *ω*^*2k*^ *P(Θ***^*2k*^**) /∑_k_ *ω*^*1k*^ *P(Θ*^*2k*^*| Θ*^*1k*^ ), where the proportionality constant is the sum over all weights (∑_k_ *ω*^*tk*^).

**Step 3:** Step 2 is repeated *T* times (*t=3,4,…,T*) when the algorithm terminates. The last set of parameter values (***Θ*^*T*^**) kept by the algorithm represent samples from the approximate posterior distribution of the model parameters (***Θ***).

Weighted Euclidean distance function was used for calculating errors (*d(****r***,***r***_*s*_^*1k*^)); Gaussian function was used for calculating both the closeness measures (*K*_*c*_(***r***_*stk*_,***r***)) and proposal distributions (*P(Θ*^*t*^*| Θ*^*(t-*^ _*1)k*_ )). The above algorithm was parallelized in the following manner. In each step *t*, instead of sampling one set of parameter values at a time and checking whether it passes the error threshold, *N*_*B*_ numbers of parameters values { *Θ***^*tk*^**, *k=1,…,N*_*B*_} were sampled at a time. Simulating and evaluating the local response matrices using each of these sampled values ({ *Θ***^*tk*^**, *k=1,…,N*_*B*_}) were performed in parallel across multiple processors. Since simulating each local response matrix requires solving an ODE model, which is computation intensive, performing several such simulations in parallel saves significant computation time.

### Parameter identifiability issues and potential remedies

An STN consisting of *N* proteins can have up to (*N*^*2*^*-N*) possible interactions excluding self-regulation. The local response matrix (***r***) of the STN provides a quantitative representation of each of these interactions. However a typical STN has far less interactions (*N*_*c*_) than theoretically possible,*N*_*c*_ **<<** (*N*^*2*^*-N*). The LRCs corresponding to the non-self-regulatory interactions that are theoretically possible but do not occur in reality are close to zero (*17*) and do not contribute in the parameter inference process. The remaining *N*_*c*_ LRCs are useful for fitting parameters. However, in a typical scenario, a mathematical model requires more than *N*_*c*_ parameters to formulate *N*_*c*_ interactions, i.e. the number of parameters (*N*_*p*_) in the model is typically larger than the number of interactions (*N*_*c*_) it formulates (*N*_*p*_ *> N*_*c*_). Generally speaking, fitting a model with less data points than the number of model parameters causes parameter identifiability and model overfitting problems. There are several ways of avoiding this problem as described below.

- One way of resolving the parameter identifiability problem is to generate SSPR data in different experimental conditions. For instance, the STN can be stimulated with different ligands or different doses of the same ligand, and following each stimulation the full set of perturbation experiments (including unperturbed measurements) needs to be performed. This will allow one to calculate multiple local response matrices (***r***^*′*^, *l=1,2,…*) for the same STN. The model can then be fitted to all local response matrices simultaneously using the same method described in the previous section.
- Network features other than the local response matrix can also be obtained from multi-conditional SSPR data and used for parameter fitting. For instance, the changes in the steady state concentrations (***x***^*′*^ = { *x*^*′*^_*i*_ *, i=1,…,N*}) of the STN components due to changes in the dose or type of ligand (*l*) can also be useful for parameter calibration. To elaborate, let the steady state concentrations of the STN components in response to ligand stimulation *l* (but no other perturbation) be denoted by ***x***^*′*^ = { *x*^*′*^_*i*_ *, i=1,…,N*}. The ratio *(**ρ**^jk^_i_*) of the concentration of node *I* at two different types or doses of ligands (*l=j,k*), i.e. ***ρ**^jk^_i_* ={ *x*^*j*^_*i*_ / *x*^*k*^_*i*_ }*, i=1,…,N*, represents the change in concentration of node *i* when the ligand or ligand concentration is changed from *j* to *k.* Therefore, these ratios *(**ρ**^jk^_i_*) quantify how different ligands influence the STN components, as opposed to local response matrices which contain information about how different nodes influence each other. Here, these ratios (*(**ρ**^jk^_i_ ,i=1,….,N*) contain information which is complementary to the local response matrices and can be augmented with these matrices (***D***_*obs*_ ={***r***^*′*^*, **ρ**^jk^_i_ ; l=1,2,…; j,k=1,2…, j≠k, i=1,2,…,N* }) to further improve parameter identifiability.
- Any additional experimental data can also be incorporated in the parameter inference process, especially if it does not incur additional computational cost. For instance, time course measurements (*x*_*i*_*(t)*) of the concentrations of any node (*i*) of the STN can also be incorporated. Incorporating time course measurements do not incur any significant additional computational cost since the model needs to be simulated once per ligand or ligand concentration, with or without such data.

## Results

### Testing algorithm using simulated data

#### Simulating perturbation response of the MAPK pathway

To test our algorithm we simulated SSPR data using a mathematical model of the ERK pathway (Fig. 1A), which is a three tiered MAPK cascade that controls cell fate (*18-20*). It comprises of three kinases, RAF, MEK and ERK. RAF is at the top of the cascade which is activated by RAS-GTP when ligands such as Epidermal Growth Factor (EGF) binds to EGF receptor on the cell surface. Activated RAF (aRAF) then activates MEK by phosphorylating it on two sites. Active MEK (aMEK) in turn activates ERK by doubly phosphorylating it. Activated RAF, MEK and ERK (aERK) are subsequently inactivated by phosphatases which de-phosphorylate them. Activated ERK (aERK) can inhibit the activation and assist in the inactivation of RAF and MEK respectively, thereby forming two negative feedback loops (*17, 21, 22*). A few simplifying assumptions were made to develop a mathematical model of this pathway. For instance, while in relality EGF activates RAF via a network of adaptor proteins and RAS-GTPs, for the purpose of modelling it was assumed that RAF is directly activated by EGF. Additionally, the activations of RAF, MEK, ERK are two stage processes involving phosphorylations of two distinct sites on these kinases. For simplicity, we combined the two stage activation process of these kinases into one stage in which the inactive form of the kinase (iRAF, iMEK, iERK) are converted into their active forms (aERK, aMEK and aERK) (*23*). Finally, the negative feedbacks from aERK to aMEK and aRAF operates via two different mechanisms. Finally, in reality the two ERK mediated negative feedback loops are mediated by two different mechanisms of inhibition of upstream kinase activities (aRAF and aMEK). However, for modelling, it was assumed that both feedback are caused by aERK mediated inactivation of aMEK and aRAF. Activation and inhibition of each kinase were formulated using Michaelis Menten functions as shown below

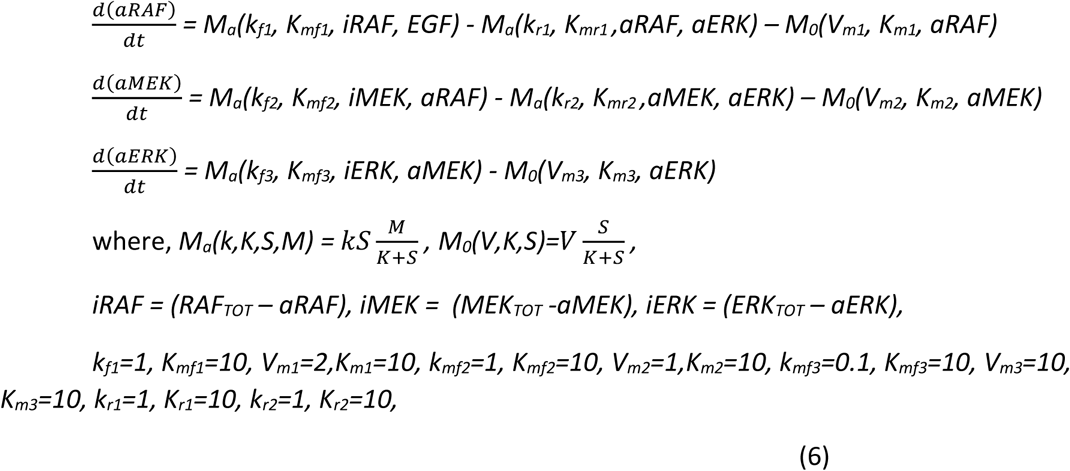

**Figure 1:**
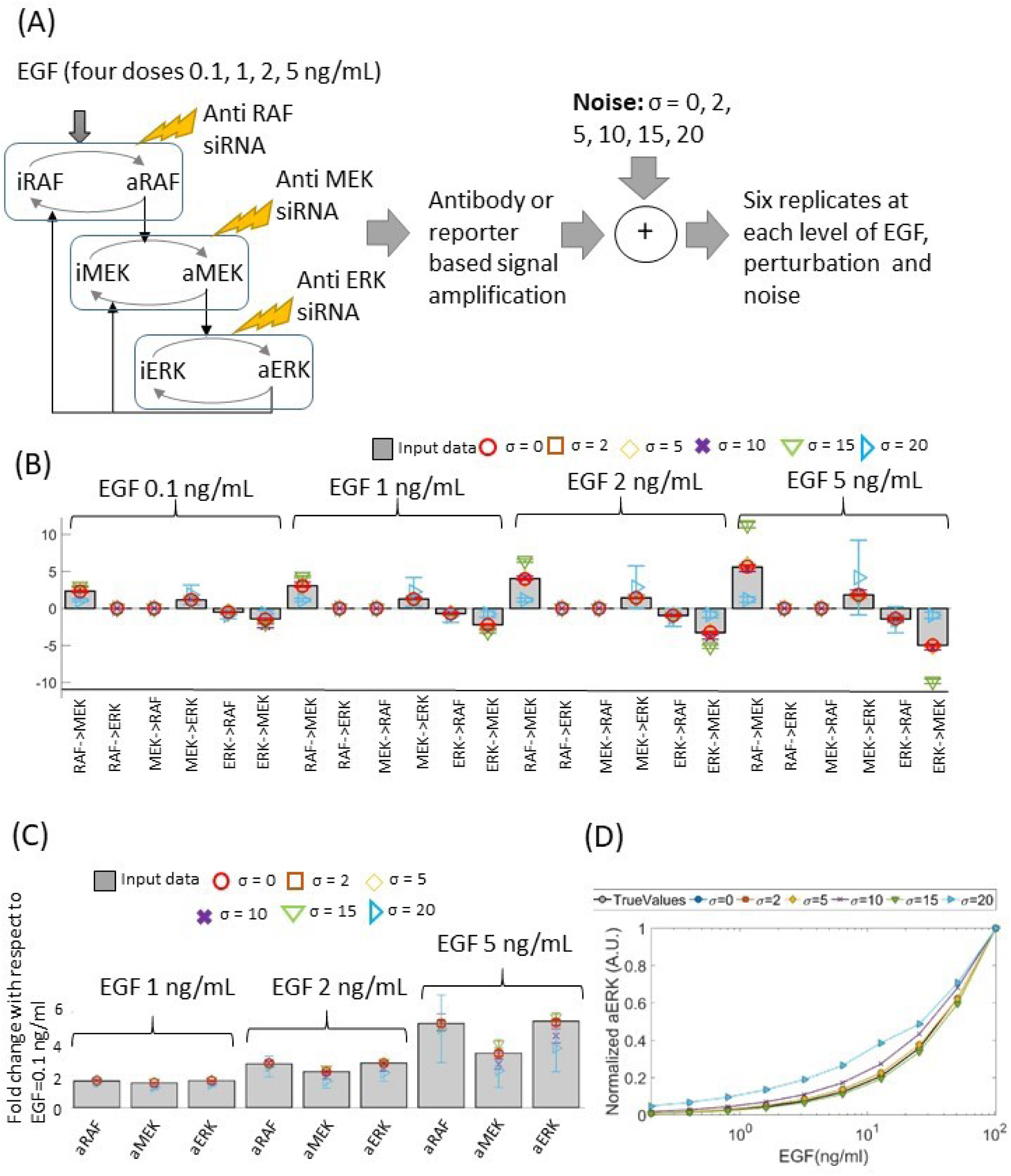
Parameter calibration using local response coefficients calculated from simulated data. (A) Schematic diagram of the MAPK model that was used to simulate perturbation response data, along with an outline of the data generation process. (B,C) Local response coefficients and steady state levels of aRAF, aMEK and aERK, simulated with the original (grey bars) and inferred parameters (coloured markers). Parameters were inferred from data contaminated with different levels (σ =0,2,5,10,15,20) of noise. The steady state levels of aRAF, aMEK and aERK at EGF levels 1,2,5 ng/mL are shown in terms of fold-change with respect to the same at EGF=0.1 ng/mL. (D) aERK levels following stimulation by different doses of EGFs.

The following experimental scenario was simulated using the above mathematical model (Eq. 6). Following common practice (*3, 23-25*), it was assumed that the cells are starved prior to stimulation by EGF. Since the phosphorylation levels of kinases are negligible in starved cells, the initial concentrations (*aRAF*_*t=0*_*, aMEK*_*t=0,*_ *aERK*_*t=0*_) of aRAF, aMEK and aERK were set to zero. The starved cells are stimulated by adding EGF to the growth medium. This was simulated by setting the EGF level of the model to a positive constant. The cells are then allowed to relax until they reach steady state. This was simulated by running the model until steady state. Once the cells attained steady state, the concentrations of active RAF, MEK and ERK are measured using antibodies or fluorescent reporters which amplify the changes in concentrations by several orders of magnitude. The amplifying effects of antibodies and reporters were simulated by multiplying the simulated concentrations by a large constant (*k*_*f*_*>>1*) (*26*). Biological measurements are typically noisy, which was simulated by adding random Gaussian noise to the amplified concentrations. Since biological data are typically generated in replicates, we generated six replicates for each measurements, each of which is a noisy realization of the amplified concentrations.

The above data represents active RAF, MEK and ERK in EGF stimulated, but otherwise unperturbed cells. To simulate perturbation experiments, it was assumed that the ERK pathway was perturbed by transfecting the cells with siRNAs targeting RAF, MEK, and ERK. Since siRNAs reduce the total concentration of their target proteins, the perturbations were simulated by reducing the total amount of RAF, MEK and ERK (aRAF+iRAF, aMEK+iMEK, aERK+iERK respectively) in the ODE model. Following each perturbation, the model was simulated until steady state, the steady state concentrations were amplified and measurement noise were added as described in the previous paragraph. Six replicate measurements were generated following each perturbation. These simulated concentrations of the aRAF, aMEK and aERK can be used to calculate the local response matrix of the ERK STN using Eqs. 4 and 5. However, since the STN has three active components, the local response matrix is a *3 X 3* matrix whose diagonal elements are by definition -1 (see. Eqs 2,3) regardless of the parameter values, leaving us with six LRCs, only four of which represent true interactions, to fit the model (Eq. 6) which has sixteen parameters. Since the model has significantly more parameters than the number of LRCs, it is evident the parameters of the model are not identifiable from a single local response matrix. To solve the model identifiability issue, we generated data for four different levels of EGF (*0.1 ng/ml, 1ng/ml, 2ng/ml, 5ng/ml*; see Fig. 1A). To evaluate the robustness of our algorithm against experimental noise, we generated data for six different levels (standard deviation *σ= 0, 2, 5, 10, 15, 20*) of noise (Fig 1A). Six replicate datasets were generated at each levels of EGF and noise (Fig. 1A).

#### Calibrating model parameters using simulated SSPR data

The ODE (Eq. 6) was separately fitted to data containing different levels of noise. At each level of noise (*σ>0*) and EGF stimulation, the means of the steady state concentrations of aRAF, aMEK and aERK were first estimated by calculating sample mean of the replicate measurements. The mean concentrations of the perturbed and the unperturbed STNs were then used to calculate four global response matrices (***R***^*E*^, *E=0.1, 1, 2, 5 ng/ml*), one for each EGF level, using Eq. 5. The global response matrices were then converted into local response matrices (***r*^*E*^**, *E=0.1, 1, 2, 5 ng/ml*) using Eq. 4. The ratios (***ρ***_*E,0.1*_*, E=1,2,5 ng/ml*) between the concentrations of aRAF, aMEK and aERK at EGF levels *1, 2, 5 ng/ml* to those at the lowest EGF level (*0.1 ng/ml*) were also calculated and were used for model fitting.

Adaptive weight ABC-SMC algorithm (*16*) was then used to calibrate the model parameters. Total concentrations of RAF, MEK and ERK were assumed to be known and therefore set to the same values that were used for data simulation. The initial concentrations of the model were set to zero reflecting characteristics of starved cells. The prior distributions of remaining sixteen model parameters was set to log-normal distributions with mean and standard deviations *2.3* and *2* respectively. For each set of parameter values, four local response matrices (***r*_*M*_^*E*^,** *E= 0.1, 1, 2, 5 ng/ml*), one for each EGF level, were calculated using Eqs. 1-3. Three sets of ligand response ratios *(**ρ***_***M***_^*E,0.1*^*, E=1,2,5 ng/ml*) were also calculated. These were then compared with those (***r*^*E*^**, *E= 0.1, 1, 2, 5 ng/ml; **ρ***^*E,0.1*^*, E=1,2,5 ng/ml*) calculated from simulated data using weighted Euclidean distance. The overall distance (*d*_*o*_) between four pairs of local response matrices (***r*^*E*^*, r*_*ME*_,** *E= 0.1, 1, 2, 5 ng/ml*) and three pairs of ligand response coefficients (***ρ***^*E,0.1*^*, **ρ***_***M***_^*E,0.1*^*, E=1,2,5 ng/ml*) was calculated as

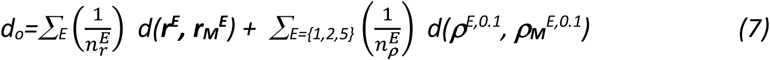

where 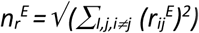 and 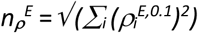 are weights, *d(****x,y***) represents Euclidian distances between **x** and **y**. At each stage (*t*) of the weighted ABC-SMC algorithm, *N*_*ABC*_=1000 sets of parameters were selected, for which the distance *d*_*o*_ is less than the error threshold *ε*_*t*_. The set of parameters (***Θ*^*T*^=**{ ***Θ*^*T*^_*i*_**, *i=1,…,N*_*abc*_}) that were selected at the final stage (*t=T*) of the algorithm were then used as samples from the posterior distributions of the parameters. Parameters were inferred separately from SSPR data sets containing different levels of noise. For national convenience, parameters sampled from data containing different levels of noise will be denoted by ***Θ*^*T*^_*σ*_** hereafter. To see whether the sampled parameters provide a good fit to the data the LRCs and the ligand response ratios of the pathway were simulated from each set of sampled parameters (***Θ*^*T*^_*σ*_,** *σ*=0,2,5,10,15,20). The mean and standard errors of these quantities are shown using markers and error bars in Fig. 1B,C. Those calculated from noise-free SSPR data are also shown in these figures using bar charts. These figures suggest that the LRCs and ligand response ratios calculated from the noise free SSPR data and simulated using the sampled parameters match closely when the noise in the data is less than σ=10. The model fit worsens at higher noise level.

#### Predicting active ERK levels in response to different doses of EGF using the calibrated models

Depending on several extrinsic and intrinsic factors such as types and concentrations of ligands, reaction rates etc. a biochemical pathways may take very different temporal trajectories to arrive at the same or very similar steady states (*27-29*). Therefore, pathway models that are fitted to steady state data can only be expected to predict the steady state behaviour of the pathway but not its kinetic behaviour. To see if the calibrated models can predict the steady state behaviour of the in-silico MAPK pathway, we simulated steady state dose response of aERK using the parameters sampled at each noise level. We chose EGF doses (EGFs=0.2 0.4 0.8 1.6 3.2 6.4 12.8 25.6 51.2 102.4]; which were not used to simulate the training data. For each noise level (*σ*=0,2,5,10,15,20), aERK levels in response to different doses of EGFs were simulated using the sampled parameter values. The means and standard errors of aERK at different EGF doses were computed and plotted for each level of noise. The gold-standard in-silico aERK dose response was also plotted in the same diagram for comparison. The simulations by calibrated models (models fitted with sampled paramters) closely matched the gold-standard data when the parameters were inferred from less noisy data (σ<20). The predictions were worse at the highest level of noise (σ=20).

#### Influence of the prior parameters on model fitting

We further investigated how the choice of the prior distribution influence parameter inference. In the simulation study discussed above we chose log normal prior distributions with mean and standard deviations *2.3* and *2* respectively for all parameters. We varied the means of the prior distributions between 1 and 1000 (mean=1, 5, 10, 20, 30, 50, 100, 1000) and for each prior mean we inferred parameters from SSPR data containing four levels of noises (σ=0,2,5,10). The inferred parameters were then used to estimate the mean local and legand response coefficients of the pathway, and the sum of squared (SSQ) distances between the estimated coefficients and the original data were calculates. The SSQs represent the model fitting errors for different prior means. The SSQs for different values of prior means at different noise levels are shown in Fig. 2. When the noise is low (σ=0,2,5), the SSQs vary between 0.2-0.4 independently of the value of the prior mean. At higher noise (σ=10), the SSQ vary between 4-6 independently of the prior mean. These results suggests the model fitting error is negligible when noise is small, it depends only on the level of noise in data and not the choice of prior mean.Therefore, the proposed algorithm is robust against choices of prior parameters.

**Figure 2:**
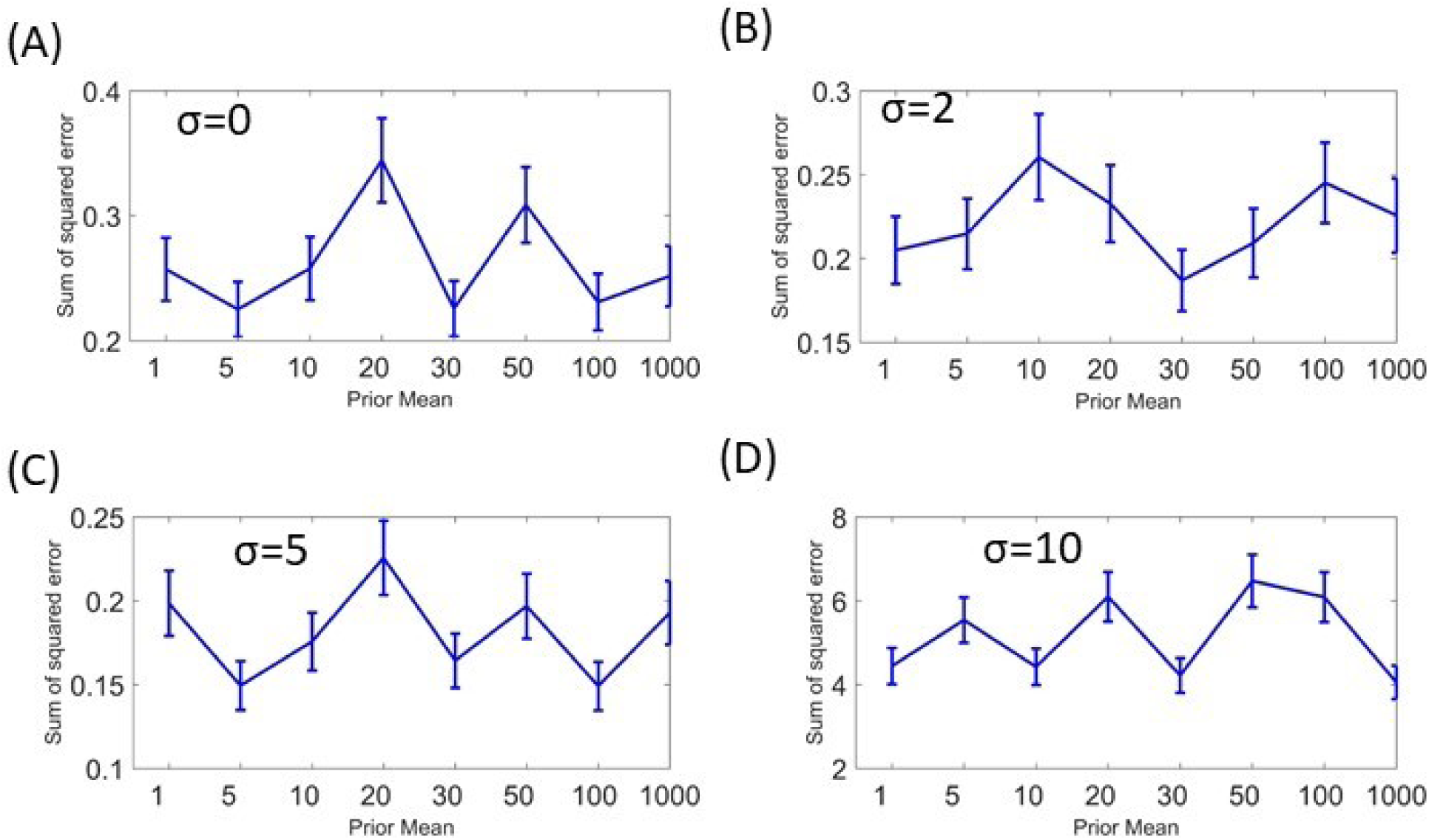
Effects of hyper-parameters on model fitting error. X-axis represents hyper-parameter values, Y-axis represent sum of square error between original and predicted LRCs and SSFCs. Error bars represent standanrd-deviations. Panels (A-D) show the effect of hyper-parameter choice on the ABPIPRD algorithm at different levels of measurement noise (σ=0,2,5,10 respectively).

### Implementing the proposed algorithm on real SSPR data

#### Calibrating an ODE model of the MAPK pathway using experimental data

We further implemented the algorithm on a real SSPR dataset that was generated to study an interesting biological phenomena involving PC12 cells which are derived from pheochromocytoma of the rat adrenal medulla. These cells proliferate and differentiate when stimulated by EGF and Nerve Growth Factor (NGF) respectively despite the fact that both of these ligands activate the ERK pathway via the same receptor (the EGFR receptor). The molecular mechanism by which EGF and NGF, which activate the same pathway, induces two different phenotypes is a matter of continued research. Santos et. al. experimentally perturbed the ERK pathway in EGF and NGF stimulated cells to understand how these ligands induce different phenotypes via the same pathway (*25*). The experiments involved treating PC12 cells by EGF or NGF, after perturbing the components (RAF, MEK, and ERK) of the ERK pathway using siRNAs. The phosphorylation levels of RAF, MEK and ERK were measured at pseudo steady-states following each perturbation and stimulation. The same were also measured without any perturbation. Using the resulting data, Santos et. al. calculated the LRCs for EGF and NGF stimulated ERK pathway (Fig. 3A)(*25*). The LRCs in response to EGF and NGF indicate that the ERK pathways might have different topologies depending on which ligand was used to stimulate it (Fig. 3A). When EGF was used, the interaction from ERK to RAF had a negative LRC indicating the presence of a negative feedback in this condition (Fig. 3A). But when NGF was used, the LRC of the same interaction was positive, indicating the presence of a positive feedback loop. Additionally, the interaction from RAF to ERK had a relatively high positive LRC when the cells were treated with NGF, but this LRC was negligibly small in the presence of EGF. Therefore, it was concluded by Santos et. al. that there was a feedforward loop from RAF to ERK in presence of NGF, but this loop was not operational in presence of EGF (Fig. 3A). To test our algorithm on Santos et. al.’s dataset, an ODE model which accounted for the topological variations of the ERK pathway in response to EGF and NGF stimulations was developed. The model is shown below.

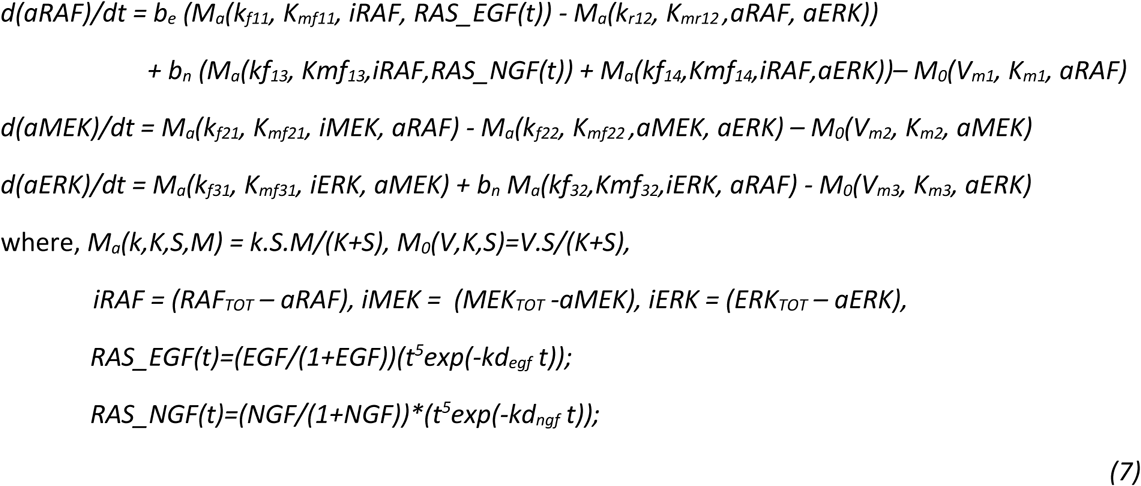

**Figure 3:**
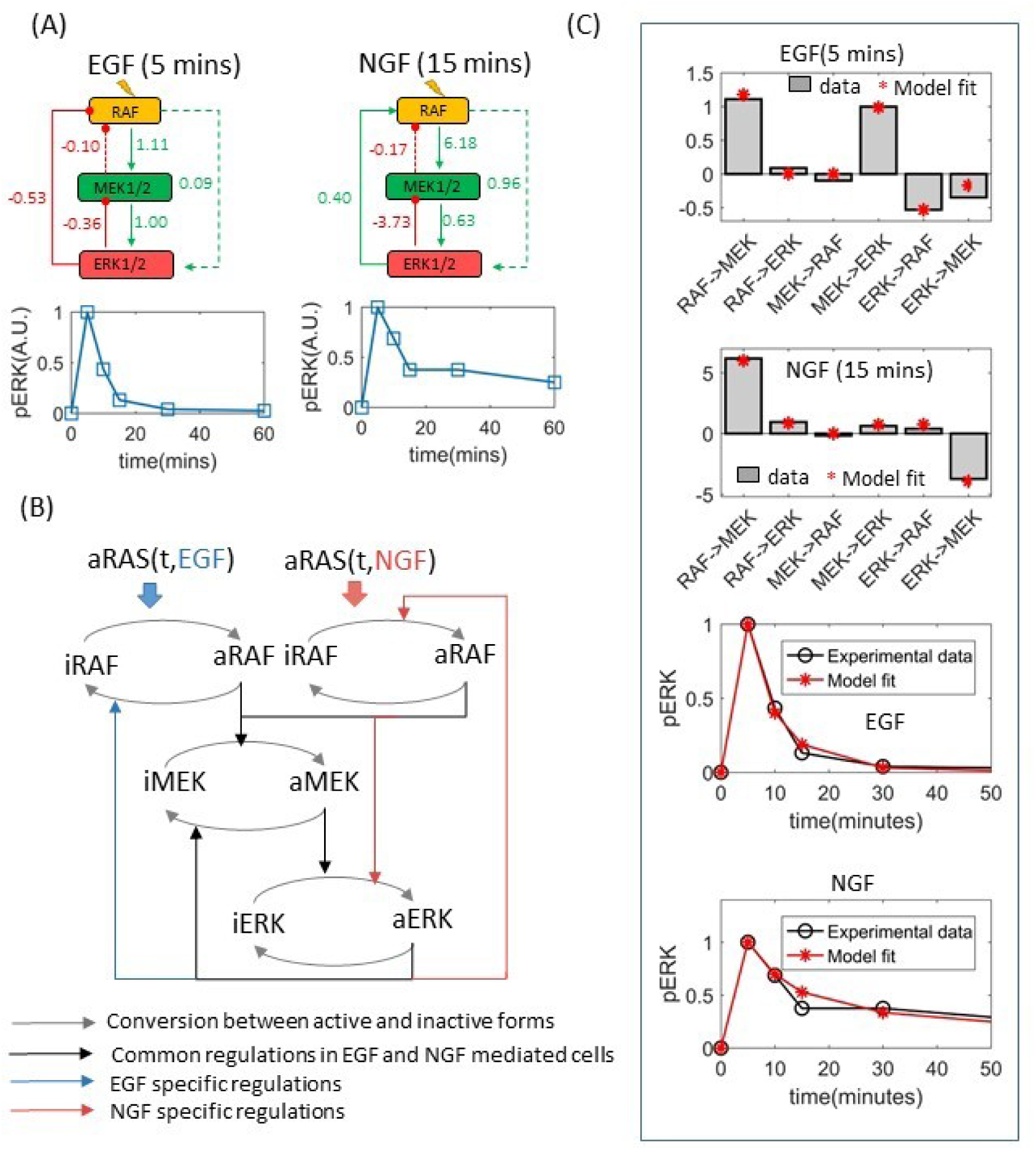
Fitting an ODE model of the MAPK pathway to experimental data. (A) LRCs of the ERK pathway and time-dependent relative pERK concentrations in EGF and NGF stimulated PC12 cells. (B) Schematic diagram of the ODE model that was fitted to the data presented in (A). (C) LRCs and time dependent pERK concentrations calculated using the fitted models. LRCs calculated from experimental data and experimentally observed pERK kinetics are also shown in this panel for comparison. Model fits represent average of an ensemble of one thousand models fitted to 1000 sets of parameters sampled by the variable weight ABC-SMC algorithm. Error bars represent standard error. Error bars are not visible due to having negligible standard error.

The above model (Eq. 7) of the ERK pathway is in some ways different from the one (Eq. 6) used for the simulation study.

- Firstly, unlike in the previous case (Eq. 6), it is no longer assumed that EGF or NGF directly activates RAF. This simplification step is avoided to reflect the biological reality that RAF is activated by the RAS proteins which are activated by EGF and NGF via a series of biochemical interactions involving the receptor and adaptor proteins. The SSPR dataset does not encompass receptor, adaptor and RAS proteins, therefore these are not exclusively incorporated in the above model (Eq. 7). But, it is known that RAS, which directly activates RAF, experiences rapid activation and successive deactivation following ligand (EGF, NGF) stimulation (*23*). This transient nature of the input signal to RAF was formulated using gamma functions (*RAS_EGF(t), RAS_NGF(t)*) with unknown parameters (*kd*_*egf*_*, kd*_*ngf*_) which were inferred from data.
- Secondly, the model in Eq.7 incorporates the influence of NGF on the kinetics and the topology of the ERK pathway. Since EGF and NGF activate RAF via RAS at different rates, the influence of these ligands on RAF were formulated using two separate Michelis Menten functions (Eq. 7, Fig. 3B). Binary variables *b*_*e*_ and *b*_*n*_ were used to characterize interactions which occur selectively in response to EGF and NGF respectively (Fig. 3B).

The resulting model has twenty four unknown parameters. Additionally, the total concentrations (*RAF*_*TOT*_, *MEK*_*TOT*_, *ERK*_*TOT*_) of RAF, MEK and ERK are also unknown and therefore need to be estimated. However, there are only two sets of LRCs, a total of 12 data points (excluding LRCs for self interactions which are by definition -1) available to fit the model. Fitting such a parameter rich model using such a small number of data points will almost certainly run into model identification problems. Additional data was incorporated in the inference process. Santos et. al. measured the SSPR data at 5 and 15 minutes after EGF and NGF stimulation respectively since the PC12 cells were seen to reach pseudo steady statse at these time points. This implies that the rates of changes (*d(aRAF)/dt, d(aMEK)/dt, d(aERK)/dt*) in the phosphorylation levels temporarily became zero (*d(aRAF)/dt = 0, d(aMEK)/dt = 0, d(aERK)/dt = 0*) at these time points. This provides us six additional data points, i.e. the rates of changes in aRAF, aMEK, and aERK at 5 and 15 minutes after EGF and NGF stimulation respectively, totaling 18 data points which is still too little to calibrate a model with 27 parameters. Therefore, phosphorylation levels of ERK measured at 0, 5, 10, 15, 30 and 60 minutes following EGF and NGF stimulation (*25*) were also incorporated in our inference algorithm to supplement the LRCs and pseudo steady state data.

For parameter inference it was assumed that all model parameters and the total concentrations RAF_TOT_, MEK_TOT_ and ERK_TOT_ have log-normal prior distributions. The means of the prior distributions of all model parameters except those of the gamma functions (*kd*_*egf*_*, kd*_*ngf*_) were set to 2, those of the gamma function parameters (*kd*_*egf*_*, kd*_*ngf*_) were set to 0.2, and those of the total concentrations RAF_TOT_, MEK_TOT_ and ERK_TOT_ were set to 25, 100 and 400 respectively. The standard deviations of all priors were set to 2. The initial concentrations of aRAF, aMEK and aERK were all set to 0 since in Santos et. al’s experiments cell were starved prior to stimulations. The Weighted ABC-SMC based algorithm was run using the above settings. LRCs of the ERK pathway model (Eq.7) and temporal activities of aERK in response to EGF and NGF were simulated using the inferred parameters and then plotted against those derived from experimental data (Fig. 3C), showing a close match between the two. The inferred parameters were then used to predict different kinetic and steady state pathway behaviours which were not used for model calibration.

#### Predicting active RAF and MEK levels in EGF and NGF stimulated PC12 cells using the calibrated model

Firstly, the relative changes in the concentrations of aRAF and aMEK within a 60 minutes period after EGF and NGF stimulations were simulated. Simulations suggested that aRAF and aMEK levels peak at 5 minutes after both EGF and NGF stimulations, but diminish much quicker after EGF stimulation than NGF stimulation. The simulated aRAF and aMEK activities (Fig. 4A) qualitatively reflected the experimental data (*25*).

**Figure 4:**
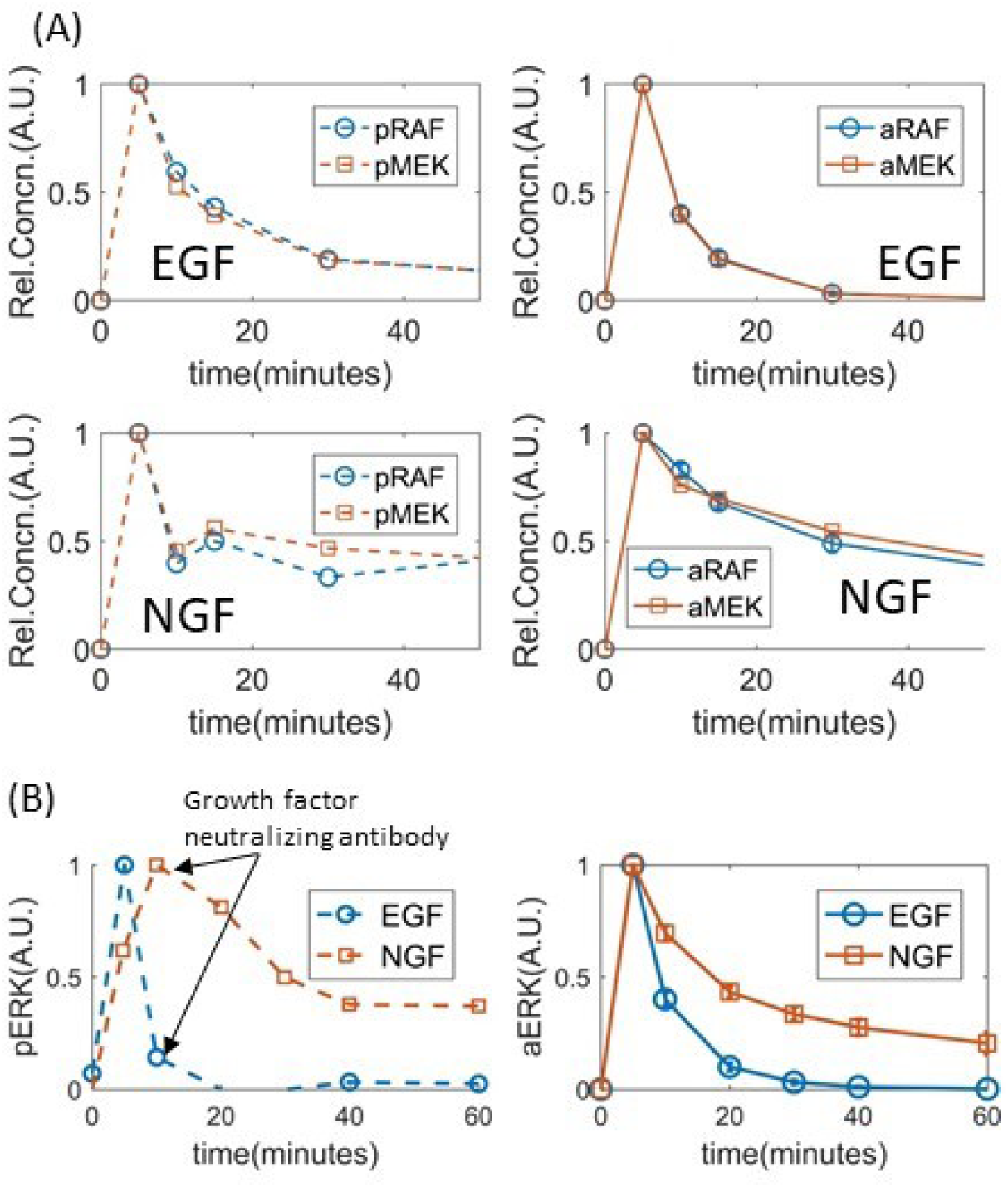
Simulating temporal concentrations of pRAF and pMEK using the fitted models. (A) Time dependent relative concentrations of pRAF and pMEK in response to EGF (top two sub-panels) and NGF (bottom two subpanels). Experimental data are shown in the left sub-panels and the model simulations are shown in the right sub-panels. (B) Temporal response of pERK to the application of growth factor neutralizing antibody at 10 minutes. The left and right sub-panels show experimental data and model simulation respectively. Model simulations represent average of an ensemble of one thousand models fitted to different sets of parameters sampled by the variable weight ABC-SMC algorithm. Error bars represent standard error.

#### Predicting the effect of growth factor neutralizing antibodies on aERK level in PC12 cells

The response of the ERK pathway to growth factor neutralizing antibodies applied at 10 minutes after EGF and NGF stimulation were simulated using the sampled parameters. The effect of the neutralizers were formulated by setting the ligand concentrations to zero after 10 minutes. The simulation results partially agreed with the experimentally observed behaviour (Fig. 4B). In simulation, aERK level diminished completely at 60 minutes after EGF stimulation; whereas following NGF stimulation aERK level diminished at ∼22% of its peak value at the same time point (Fig. 4B).While these general trends were also observed in experimental data (obtained from (*25*) and also shown in Fig. 4 for convenience), there were also some differences between the simulation and experimental data. The first noticeable difference is that aERK level diminished significantly faster between its peak at 5 minutes and 10 minutes (when the growth factor neutralizing antibody was applied) after EGF stimulation in the experimental observations, compared to the simulation (Fig. 4B). The second obvious difference is that the aERK level peaked at 10 minutes after NGF stimulation in the biochemical experiments, but in simulation the peak occurred at 5 minutes (Fig. 4B). In both cases, the differences between the experimental data and model simulation occur before the application of growth factor neutralizing antibodies. Therefore, it is unlikely that the difference is caused by error in simulating the effect of the neutralizing factors. A closer look at the two sets of experimentally measured phospho-ERK levels, one without the neutralizers and was used for model calibration (Fig. 3C) and the other with the neutralizers (Fig. 4B), reveals that these two sets of measurements are at odds with other. This is most likely due to biological variability between the samples used in these two experiments and/or batch effects. Therefore, in this case, the apparent differences in experimental data and model simulation can be attributed to these factors.

#### Predicting active ERK concentrations in response to different doses of EGF and NGF

The response of aERK at five minutes following different doses of EGFs and NGFs were simulated (Fig.5A) using the sampled parameters. The average simulated aERK levels in response to different doses of EGF and NGF are shown in Fig. 5A. Two different sigmoidal curves were fitted to the EGF and NGF dose responses of aERK, mainly to show that (at 5 minutes after stimulation) the concentration of active ERK has a sigmoidal relationship with those of these ligands (5A). Similar sigmoidal relationship between ligand concentrations and active-ERK levels (at five minutes after stimulation) were experimentally observed in PC12 cells (data obtained from (*30*), also shown in Fig. 5A for convenience).

**Figure 5:**
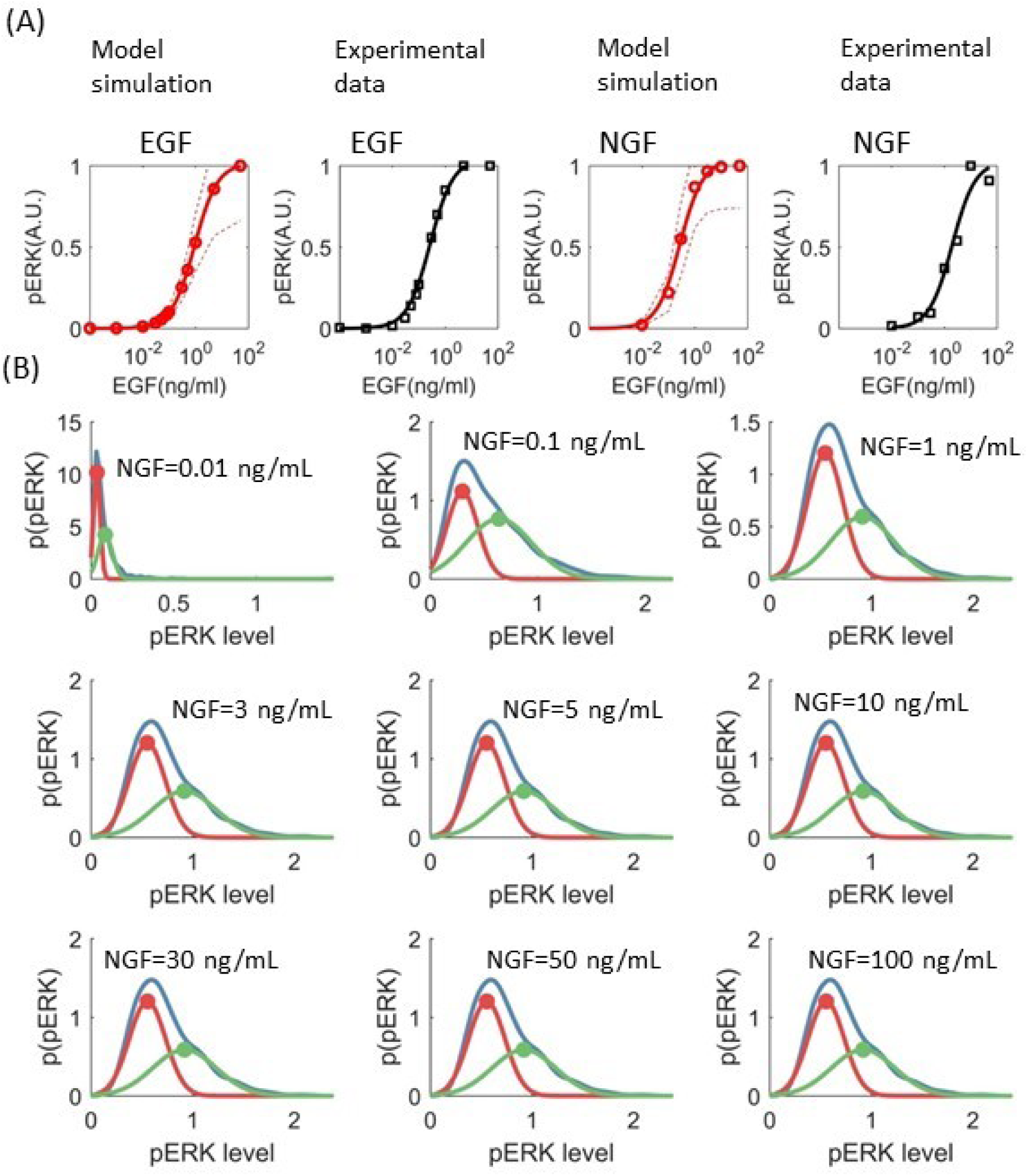
pERK concentrations at different doses of growth factors. (A) Simulated (shown in red) and experimentally measured (shown in black) relative pERK concentrations following five minutes of EGF and NGF treatments. A.U. means arbitrary units. The dashed lines represent 67% confidence interval. An ensemble of one thousand models fitted to different sets of parameters sampled by the VW-ABC-SMC algorithm were used to calculate mean response (solid red lines in panel A) and confidence intervals (dashed red lines in panel A). Bimodal distribution of steady-state (60 minutes after NGF stimulation) pERK levels following treatment by different doses of NGF. For each level of NGF, pERK levels were simulated using an ensemble of a thousand models. The empirical distributions (the blue lines in panel B) of the simulated pERK levels are shown in blow. Individual Gaussian components that make up the empirical distributions are shown in red and green. The peak of the individual components are marked using dots of the respective colour.

#### Predicting active ERK concentrations in response to different doses of NGF at single cell resolutoin

The inferred parameters were used to simulate steady state response of aERK to different doses of NGF at a single cell level. To do so, each sampled set of parameter values were assumed to represent a single cell, thereby, the ensemble of all sampled parameter sets represents a cell population. Steady state aERK levels (at 60 minutes) were simulated for each set of parameter values at each level of NGF (0.01, 0.1, 1, 3, 5, 10, 30, 50, 100 ng/ml). The distribution of steady state aERK levels in a cell population in response to different doses of NGF were estimated using a kernel density estimator (https://uk.mathworks.com/help/stats/ksdensity.html). It was previously shown that at steady state, in response to NGF > 1ng/ml phosphorylated ERK levels have bimodal distributions in populations of PC12 cells (*25*). To see if the same is true for the simulated aERK levels, we fitted one or two Gaussian probability density functions to each of the aERK distributions depending on whichever produced the minimum fitting (SSQ) error. In all cases, two Gaussian Distributions provided better fits than a single Gaussian distribution, suggesting that, in our simulations, aERK has bimodal distribution at all levels (0.01, 0.1, 1, 3, 5, 10, 30, 50, 100 ng/ml) of NGF stimulation (Fig. 5B). However the locations of the different modes are nearly inseparable at NGF=0.01 ng/ml and have the highest separations at or more than 1ng/ml NGF. Therefore, the overall simulation results largely reflects the experimental observation with one exception which occur at NGF = 0.1 ng/ml. At this concentration of NGF the simulated aERK levels were seen to have bimodal distribution whereas experimentally observed phospho-ERK levels had a single mode. One possible reason behind this difference is that, in experiments, the phospho-ERK levels were measured at 16 hours after EGF stimulation. By that time, the ERK pathway is known to be influence by transcriptional events which are not accounted for in our model (*31-33*). This might cause some differences between the dose responses of the model and the real ERK pathway.

## Discussion

We developed an algorithm that can calibrate ODE models to SSPR data without exclusively simulating the perturbation experiments during the calibration process. This has several benefits beyond reducing computational cost. For instance, in many scenarios exact mechanism or ‘direct’effect of the biochemical perturbations are not known, making it impossible to simulate these experiments in the first place. For example, the mechanism of action or the exact targets of biochemical inhibitors are often either not known or not straightforward to incorporate in a model without significantly increasing the model complexity. Therefore, the data produced by the perturbation experiments where such inhibitors are used are not useful for fitting ODE models. Our approach does not require any knowledge of the perturbation experiments except which protein is being perturbed, thereby expanding the periphery of usable data for fitting ODE models. The models fitted using our algorithm were shown to be able to largely reproduce STN behavior both at population and single cell level. However, this approach of model fitting is not without its caveats. It relies on fitting parameters of a model to the LRCs of the STN. In any condition, a STN only has as many LRCs as the number of its interactions. Since each of these interactions are formulated using kinetic equations that usually have more than one parameters, in almost all cases there are more parameters to fit than the number available LRCs. However, the behavior of biochemical networks varies depending on the dose and type of ligand stimulations, and so do the LRCs of the systems.

Therefore, it is possible to estimate LRCs of the STN in response to different doses or types of ligand stimulations and use these LRCs to calibrate model. The upside of performing perturbation experiments in multiple conditions is that the resulting data is more informative than data from only one condition, but downside is the increased experimental burden.

Another potential weakness of our method is that it inherently relies on the LRCs, and therefore on the accuracies of the algorithms, that are used to calculate LRCs from experimental data. Several such algorithms had been developed for this purpose. The classical MRA algorithm (*17*) requires as many perturbations as the number of components in the network to calculate LRCs, this is experimentally tedious and sometimes not feasible due to lack of appropriate antibody, inhibitors etc. It also works best on noise free data, which is not feasible to produce using biochemical experiments. Many statistical reformulation of the classical MRA have since been proposed to deal with these issues (*1, 25, 31, 34, 35*). Some of these algorithms can calculate LRCs from noisy data obtained from a relatively small numbers of perturbations. How a combination of these statistical realizations of MRA and the parameter calibration algorithm proposed in this paper can be used to reconstruct and calibrate mathematical models of large signaling STNs is yet to be investigated and is a matter of future research.

## Acknowledgement

This project was funded by the Irish Cancer Society CCRC BREAST-PREDICT [grant number CCRC13GAL]

